# Antibiotic-induced dysbiosis predicts mortality in an animal model of *Clostridium difficile* infection

**DOI:** 10.1101/315382

**Authors:** Charles Burdet, Sakina Sayah-Jeanne, Thu Thuy Nguyen, Perrine Hugon, Frédérique Sablier-Gallis, Nathalie Saint-Lu, Tanguy Corbel, Stéphanie Ferreira, Mark Pulse, William Weiss, Antoine Andremont, France Mentré, Jean de Gunzburg

**Affiliations:** INSERM & Paris Diderot University, IAME, UMR 1137, Paris, France; Department of Epidemiology, Biostatistic and Clinical Research, Bichat Hospital, AP-HP, Paris, France; Da Volterra, Paris, France; GenoScreen, Lille, France; UNT Health Science Center, Fort Worth, TX 76107, USA; Laboratory of Bacteriology, Bichat Hospital, AP-HP, Paris, France

**Author notes:** These authors contributed equally to the work (France Mentré, Jean de Gunzburg). **Corresponding author:** Jean de Gunzburg, Address: Da Volterra, Le Dorian Bât B1, 172 rue de Charonne, 75011 Paris, France, Mail. **Author contributions** SSJ, FSG, NSL, TC, AA and JG designed the research. MP, WW and FSG performed the research. SF performed the metagenomic analysis. CB, TTN and FM performed the statistical analysis of the data. CB, SSJ, FM, AA and JG wrote the paper. All authors agreed on the final version of the manuscript.

**Keywords:** antibiotics, dysbiosis, *C. difficile* infection, hamster animal model, mortality, prevention

## Abstract

**Background:** Antibiotic disruption of the intestinal microbiota favors colonization by *Clostridium difficile*. Using a charcoal-based adsorbent to decrease intestinal antibiotic concentrations, we studied the relationship between antibiotic concentrations in feces and the intensity of dysbiosis, and quantified the link between this intensity and mortality.

**Methods:** We administered either moxifloxacin (n=70) or clindamycin (n=60) to hamsters by subcutaneous injection from day 1 (D_1_) to D_5_, and challenged them with a *C. difficile* toxigenic strain at D_3_. Hamsters received various doses of a charcoal-based adsorbent, DAV131A, to modulate intestinal antibiotic concentrations. Gut dysbiosis was evaluated at D_0_ and D_3_ using diversity indices determined from 16S rRNA gene profiling. Survival was monitored until D_16_. We analyzed the relationship between fecal antibiotic concentrations and dysbiosis at the time of *C. difficile* challenge and studied their capacity to predict subsequent death of the animals.

**Results:** Increasing doses of DAV131A reduced fecal concentrations of both antibiotics, lowered dysbiosis and increased survival from 0% to 100%. Mortality was related to the level of dysbiosis (p<10^−5^ for the change of Shannon index in moxifloxacin-treated animals and p<10^−9^ in clindamycin-treated animals). The Shannon diversity index and unweighted UniFrac distance best predicted death, with areas under the ROC curve of 0.89 [95%CI, 0.82;0.95] and 0.95 [0.90;0.98], respectively.

**Conclusions:** Altogether, moxifloxacin and clindamycin disrupted the diversity of the intestinal microbiota with a dependency to the DAV131A dose; mortality after *C. difficile* challenge was related to the intensity of dysbiosis in a similar manner with the two antibiotics.

## Introduction

Antibiotics disrupt the structure and composition of the intestinal microbiota, and alter metabolic processes occurring in the gut with possible acute and long-term consequences (1-5). Short-term effects include diarrhea in 5-25% of antibiotic-treated patients, and antibiotics are the main risk factor of *C. difficile* infection (6), which causes a wide range of symptoms from mild diarrhea to toxic megacolon with an annual mortality estimated to 29,000 deaths in the United States (7, 8). The lincosamide antibiotic clindamycin, as well as fluoroquinolones are among the main antibiotic classes associated with *C. difficile* infection (9).

The burden of *C. difficile* infection increases (10), and *C. difficile* is considered by the US CDC as an urgent threat (11). *C. difficile* pathophysiology is related to the perturbation of the intestinal microbiota and its metabolism, which allows *C. difficile* spores to germinate and colonize the gut, and cytotoxic toxins to be released. Various animal models have been developed to delineate the pathophysiology of *C. difficile* infection (12); including the golden Syrian hamster model (13). In this model, hamsters treated with antibiotics and colonized by *C. difficile* are highly susceptible to lethal infection, and the degree of susceptibility to develop infection varies between classes of antibiotics (14, 15).

There is however no precise and quantitative analysis of the relationship between antibiotics effect on the global bacterial diversity within the intestinal microbiota and the development of severe *C. difficile* infection. Yet diversity is the first descriptor of the structure of a community and is believed to be a major determinant of its dynamics. The analysis of complex bacterial communities was made possible by the development of efficient sequencing technologies applied to 16S rRNA genes (16). These genes are found in all bacterial species and contain regions which are highly conserved and others which are highly variable in sequence and can be used as molecular fingerprints. Several metrics are available for measuring diversity in bacterial communities. Alpha-diversity refers to within-sample diversity and is usually analyzed using the number (richness) and the distribution (evenness) of bacterial taxa observed within a single population, e.g., the Shannon diversity index, number of observed operational taxonomic units (OTUs) and the Chao1 index (17, 18). Beta-diversity, which refers to diversity between samples, measures the distance between pairs of samples, e.g., UniFrac distances based on bacterial taxonomy and Bray-Curtis dissimilarity index (19, 20).

However whether the intensity of the dysbiosis, as it can be reflected by the variations of these global indices is quantitatively related to the occurrence of *C. difficile* infection has not be explored so far. This is important for a better understanding of the pathophysiology of *C. difficile* infection and to determine whether various degrees of dysbiosis are associated with various degrees of risk of *C. difficile* infection. Here we explored this relationship in an animal model of *C. difficile* infection in hamsters. We had previously showed that DAV131A, a charcoal-based adsorbent with the same principle of action than the DAV132 product which has recently proven to be highly effective to reduce fecal antibiotic concentrations and dysbiosis in human volunteers treated with moxifloxacin (21), reduced mortality through a decrease of fecal antibiotic concentrations in a hamster model of lethal moxifloxacin-induced *C. difficile* colitis (22). We induced various degrees of dysbiosis by treating hamsters either with clindamycin or moxifloxacin, which have different antibacterial spectra but are both highly associated with the occurrence of *C. difficile* infection, and we modulated intestinal antibiotic concentrations by using various doses of DAV131A.

## Results

In order to further analyze the pathophysiology of severe *C. difficile* colitis, we treated hamsters with either moxifloxacin (total number of 70 animals) or clindamycin (total of 60 animals) for 5 days in 2 separate studies with similar designs (see Figure 1). Some groups received various doses of DAV131A given orally *bis in die* (bid) concomitantly with the antibiotic and for an additional 3 days after (corresponding to a total of 8 days). All hamsters were challenged with 10^4^ spores of a toxigenic *C. difficile* strain on the 3^rd^ day of antibiotic treatment. We analyzed the quantitative relationship between antibiotic-induced dysbiosis at the time of the *C. difficile* challenge and the occurrence of subsequent death from infection.

**Figure 1.**
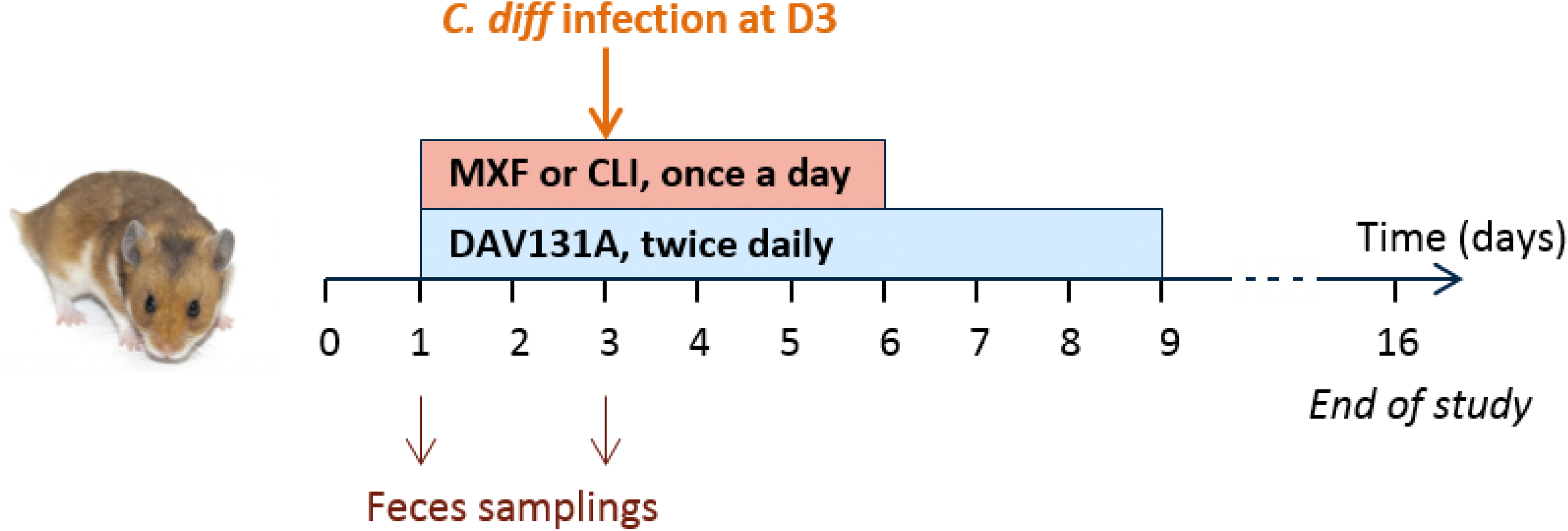
Experimental design of the studies. Male Syrian Golden hamsters were treated with moxifloxacin (MXF, n=70) or clindamycin (CLI, n=60) once a day (OAD) by the subcutaneous route for 5 days and received various doses of DAV131A *bis in die* (BID) by the oral route for 8 days, that would result in the exposition of the microbiota to various antibiotic concentrations and different bacterial environment. Toxigenic strain of *C. difficile* UNT103- 1 was inoculated at D3. Fecal samples were obtained just before the beginning of treatment, and at the 3^rd^ treatment day. Microbiota analysis was performed by 16S rRNA gene sequencing on both samples, and fecal concentration of active antibiotic was determined at D3 by microbiological assay. Survival was monitored up to D16.

Among antibiotic-treated animals, 10 (12.5%) died in the moxifloxacin study, and 28 (40.6%) in the clindamycin study. One hamster from one of the control groups died during the acclimation period, but none did after the beginning of antibiotic treatment. Significant differences in mortality rates were observed between groups which received various doses of DAV131A in addition to the antibiotic (p<10^−11^ in the moxifloxacin study and p<10^−10^ in the clindamycin study). In both studies, all hamsters treated with antibiotic + DAV131A placebo died. In the moxifloxacin study, all hamsters receiving 200 mg/kg DAV131A bid and greater survived, whereas in the clindamycin study, there was a dose-dependent reduction of mortality from 90% at 300 mg/kg DAV131 bid to 100% survival reached at 750 mg/kg DAV131 bid and above. Full results are presented in Table 1 and Figure 2.

**Table 1.** Mortality rates, fecal concentrations of active antibiotic at D3, change of Shannon index between D0 and D3 and unweighted UniFrac distances between D0 and D3 according to treatment groups in the moxifloxacin and clindamycin studies. Data are presented as n (%) or median (min; max) as appropriate. q-values refer to the comparison of the corresponding treatment group with the antibiotic + DAV131A placebo treatment group (MXF/0 or CLI/0), after Benjamini-Hochberg correction. The p-values for the comparison of all treatment groups using Fisher exact or Kruskall-Wallis tests are reported in the “All groups” line. In the analysis of concentrations, only antibiotic-treated groups were included.

| Treatment group | n | Mortality | q-value | Concentration (µg/g) | q-value | Shannon index change | q-value | Unweighted UniFrac | q-value |
| --- | --- | --- | --- | --- | --- | --- | --- | --- | --- |
| **Moxifloxacin study** |  |  |  |  |  |  |  |  |  |
| controls | 10 | 0 (0) | - | - | - | -0.1 (-0.6; 2.3) | - | 0.28 (0.24; 0.69) | - |
| MXF/0 | 10 | 10 (100) | - | 110.0 (70.5; 172.7) | - | -1.7 (-3.0; -1.0) | - | 0.61 (0.56; 0.76) | - |
| MXF/200 | 20 | 0 (0) | <10<sup>-7</sup> | 94.6 (43.0; 162.1) | 0.14 | -1 (-1.9; 0.1) | 0.00065 | 0.56 (0.49; 0.65) | 0.00075 |
| MXF/300 | 20 | 0 (0) | <10<sup>-7</sup> | 57.2 (20.2; 107.4) | <10<sup>-4</sup> | -1 (-1.5; -0.2) | 0.00026 | 0.51 (0.45; 0.57) | <10<sup>-6</sup> |
| MXF/600 | 10 | 0 (0) | <10<sup>-4</sup> | 4.2 (1.1; 20.3) | <10<sup>-4</sup> | -0.8 (-1.3; -0.3) | 0.00023 | 0.45 (0.42; 0.56) | <10<sup>-4</sup> |
| MXF/900 | 10 | 0 (0) | <10<sup>-4</sup> | 0.0 (0.0; 1.3) | 0.00033 | -0.8 (-1.6; -0.2) | 0.0022 | 0.41 (0.37; 0.51) | <10<sup>-4</sup> |
| All groups | 80 | 10 (12.5) | <10<sup>-11</sup> | 59.4 (0.0; 172.7) | <10<sup>-9</sup> | -1 (-3; 2.3) | <10<sup>-5</sup> | 0.51 (0.24; 0.76) | <10<sup>-9</sup> |
| **Clindamycin study** |  |  |  |  |  |  |  |  |  |
| controls | 9 | 0 (0) | - | - | - | 0.0 (-0.3; 0.5) | - | 0.31 (0.27; 0.38) | - |
| CLI/0 | 10 | 10 (100) | - | 10.1 (0.0; 37.8) | - | -3.9 (-4.3; -2.6) | - | 0.77 (0.72; 0.84) | - |
| CLI/300 | 10 | 9 (90) | >0.99 | 4.2 (0.0; 14.2) | 0.061 | -2.1 (-2.6; -1.5) | <10<sup>-4</sup> | 0.69 (0.62; 0.74) | <10<sup>-4</sup> |
| CLI/450 | 10 | 6 (60) | 0.11 | 0.0 (0.0; 6.5) | 0.0041 | -1.4 (-2.5; -1.0) | <10<sup>-4</sup> | 0.66 (0.63; 0.71) | <10<sup>-4</sup> |
| CLI/600 | 10 | 3 (30) | 0.0052 | 0.0 (0.0; 30.0) | 0.024 | -1.1 (-1.6; -0.4) | <10<sup>-4</sup> | 0.61 (0.55; 0.65) | <10<sup>-4</sup> |
| CLI/750 | 10 | 0 (0) | <10<sup>-4</sup> | 0.0 (0.0; 0.0) | 0.0019 | -0.8 (-1.4; 0.0) | <10<sup>-4</sup> | 0.57 (0.55; 0.61) | <10<sup>-4</sup> |
| CLI/900 | 10 | 0 (0) | <10<sup>-4</sup> | 0.0 (0.0; 0.0) | 0.0019 | -0.9 (-1.5; 0.7) | <10<sup>-4</sup> | 0.57 (0.49; 0.63) | <10<sup>-4</sup> |
| All groups | 69 | 28 (40.6) | <10<sup>-10</sup> | 0.0 (0.0; 37.8) | <10<sup>-4</sup> | -1.1 (-4.3; 0.5) | <10<sup>-9</sup> | 0.62 (0.27; 0.84) | <10<sup>-10</sup> |

**Figure 2.**
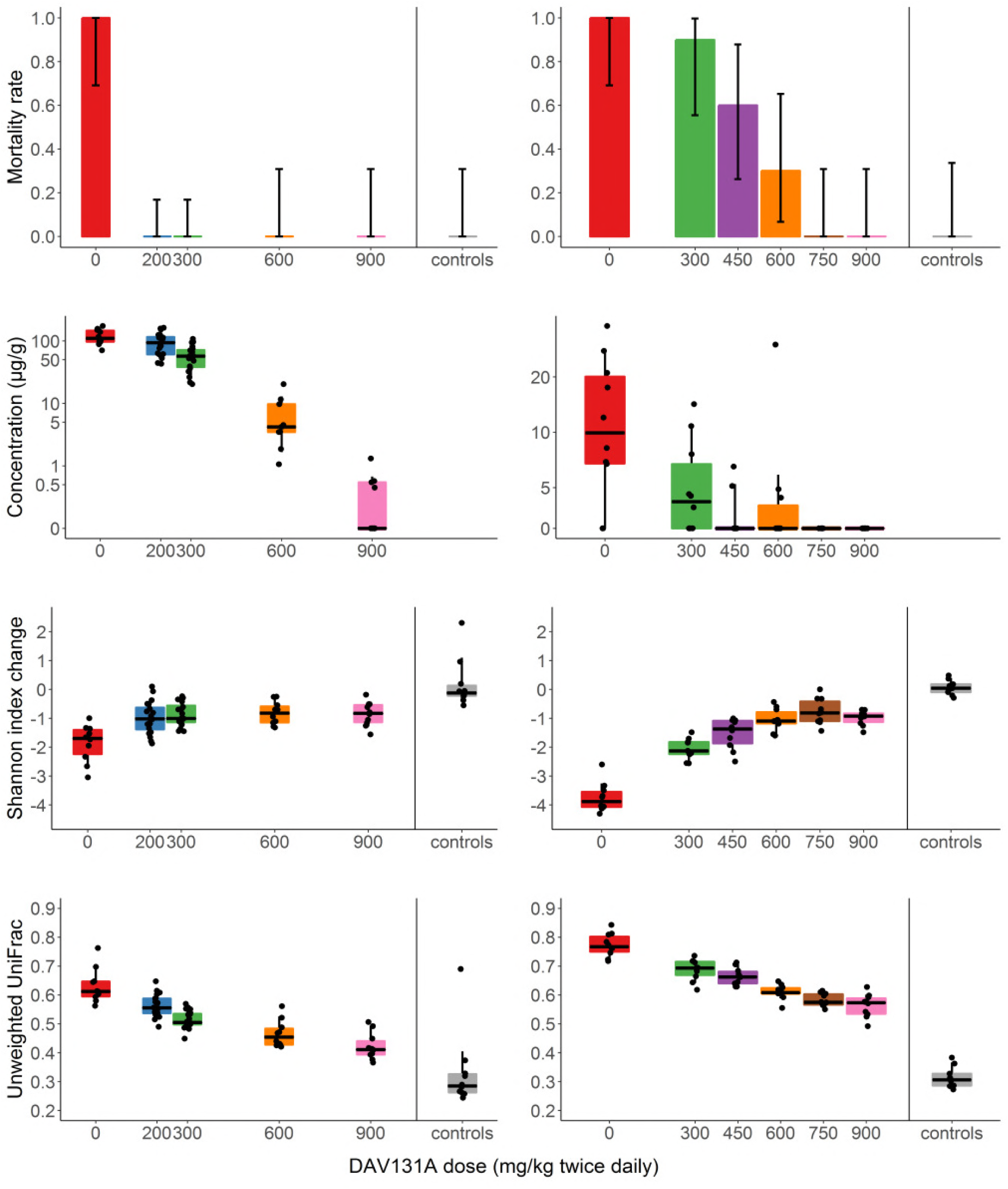
Mortality rate, fecal concentration of active antibiotic, change of Shannon index and unweighted UniFrac distance between D_0_ and D_3_ according to treatment group in the moxifloxacin (left panel) and clindamycin (right panel) studies. Barplots of the mortality rates are presented with their 95% binomial confidence intervals. For concentrations, Shannon index and unweighted UniFrac distances, the boxes present the 25^th^ and 75^th^ percentiles and the horizontal black bar report the median value, while whiskers report 5^th^ and 95^th^ percentiles.

Fecal concentration of active antibiotics, as measured by a microbiological assay, decreased as expected with increasing doses of DAV131A (p<10^−9^ in the moxifloxacin study and p<10^−4^ in the clindamycin study). These concentrations were significantly lower in hamsters which survived than in those which died during the study (p=0.00025 in the moxifloxacin study and p<10^−6^ in the clindamycin study, see Table 2).

**Table 2.** Median (min; max) values of active antibiotic concentration and change of α- (Shannon index, number of OTUs and Chao1 index) and β- (unweighted and weighted UniFrac distances, and Bray-Curtis dissimilarity) diversity indices between D_0_ and D_3_ according to vital status at D_16_ in antibiotic-treated groups for each study, and their respective area under the ROC curve (AUROC) for predicting occurrence of death by D_16_. P-values refer to non-parametric Wilcoxon test.

|  | Died | Survived | p-value | AUROC [95% CI] |
| --- | --- | --- | --- | --- |
| **Moxifloxacin study** | **N=10** | **N=60** |  |  |
| Concentration | 110.0 (70.5; 172.7) | 52.9 (0.0; 162.1) | 0.00025 | 0.87 [0.76; 0.95] |
| **α-diversity** |  |  |  |  |
| Change of Shannon index | -1.7 (-3.0; -1.0) | -1.0 (-1.9; 0.1) | $<10^{-4}$ | 0.91 [0.80; 0.98] |
| Change of Number of OTUs | -135.9 (-207.3; -52.2) | -72.9 (-201.0; 74.5) | 0.001 | 0.83 [0.67; 0.95] |
| Change of Chao1 index | -137.9 (-213.4; -46.2) | -75.5 (-229.3; 83.5) | 0.002 | 0.81 [0.64; 0.93] |
| **β-diversity** |  |  |  |  |
| Unweighted UniFrac | 0.61 (0.56; 0.76) | 0.51 (0.37; 0.65) | $<10^{-5}$ | 0.95 [0.90; 0.99] |
| Weighted UniFrac | 0.33 (0.24; 0.48) | 0.26 (0.13; 0.57) | 0.02 | 0.73 [0.58; 0.87] |
| Bray-Curtis dissimilarity | 0.78 (0.63; 0.86) | 0.60 (0.31; 0.87) | $<10^{-4}$ | 0.91 [0.81; 0.99] |
| **Clindamycin study** | **N=28** | **N=32** |  |  |
| Concentration | 5.0 (0.0; 37.8) | 0.0 (0.0; 4.4) | $<10^{-6}$ | 0.81 [0.72; 0.91] |
| **α-diversity** |  |  |  |  |
| Change of Shannon index | -2.2 (-4.3; -0.4) | -1.1 (-2.6; 0.0) | $<10^{-6}$ | 0.88 [0.78; 0.96] |
| Change of Number of OTUs | -223.9 (-344.6; -75.8) | -106.8 (-202.2; -22.0) | $<10^{-5}$ | 0.86 [0.75; 0.95] |
| Change of Chao1 index | -227.7 (-358.2; -73.8) | -110.0 (-209.0; -22.7) | $<10^{-6}$ | 0.86 [0.76; 0.95] |
| **β-diversity** |  |  |  |  |
| Unweighted UniFrac | 0.71 (0.59; 0.84) | 0.60 (0.49; 0.68) | $<10^{-10}$ | 0.94 [0.88; 0.99] |
| Weighted UniFrac | 0.42 (0.24; 0.62) | 0.30 (0.24; 0.59) | $<10^{-6}$ | 0.87 [0.76; 0.96] |
| Bray-Curtis dissimilarity | 0.86 (0.71; 0.98) | 0.70 (0.61; 0.87) | $<10^{-9}$ | 0.92 [0.84; 0.97] |

The structure of the bacterial intestinal microbiota was studied by 16S rRNA gene profiling using Illumina sequencing technology. Several α- (within sample) and β- (between samples) diversity metrics were computed for investigating the individual-specific change of diversity between the time of antibiotic initiation and the time of *C. difficile* inoculation. The loss of diversity between the beginning of antibiotic treatment and *C. difficile* inoculation was lower in DAV131A-treated hamsters than in those treated with antibiotic and DAV131A placebo (Table 1 and Supplementary Table 1). Indeed, loss of diversity increased with increasing concentrations of active antibiotic in feces, attesting of a direct relationship between antibiotic exposure of the microbiota and the extent of dysbiosis (Spearman r=−0.25, p=0.043 for the change of Shannon index between D_0_ and D_3_ and r=0.71, p<10^−10^ for unweighted UniFrac distance between D_0_ and D_3_ in the moxifloxacin study, and r=− 0.49, p<10^−4^ for the change of Shannon index between D_0_ and D_3_ and r=0.57, p<10^−5^ for unweighted UniFrac distance between D_0_ and D_3_ in the clindamycin study, see Supplementary Figure 1 and in Supplementary Table 2).

We also compared the changes in diversity within the intestinal microbiota between D_0_ and D_3_ according to the vital status at D_16_. Diversity at the time of *C. difficile* challenge was significantly less affected in hamsters which survived (Table 2 and Figure 3). In the moxifloxacin study, the median (min; max) change of the Shannon index was −1.7 (−3.0; −1.0) in hamsters which died by D_16_, versus − 1.0 (−1.9; −0.1) in those which survived (p<10^−4^). In the clindamycin study, the median (min; max) change of the Shannon index was −2.2 (−4.3; −0.4) in hamsters which died by D_16_, versus −1.1 (−2.6; 0.0) in those which survived (p<10^−7^). Interestingly, the median change of Shannon index in hamsters which died was rather similar for the 2 antibiotics, in spite of their different spectra of activity and mode of action. In order to further assess the ability of diversity indices to predict death by D_16_, we computed for each diversity index the area under the Receiver Operating Curve (AUROC), which can be interpreted as the probability that the index correctly ranks 2 randomly chosen animals. AUROCs were above 0.8 for all diversity indices studied (Table 2), attesting that they are highly predictive of the outcome (23). Each index also exhibited a similar predictability of death for both antibiotics. Changes in the Shannon index at the time of challenge had the best predictability of death by D_16_ among α-diversity indices (AUROC 0.91 [95%CI, 0.80; 0.98] for moxifloxacin and 0.88 [0.78; 0.96] for clindamycin), whereas unweighted UniFrac was the most predictive β-diversity index (AUROC 0.95 [0.90; 0.99] for moxifloxacin and 0.94 [0.88; 0.99] for clindamycin). These two indices were further studied after pooling data from the two different antibiotic treatments. Overall, data from 130 antibiotic-treated animals were available, among which 38 died by D_16_ (29.2%). Logistic models of mortality by D_16_ for both diversity indices are presented in Figure 4. The AUROC of the Shannon index change was 0.89 [0.82; 0.95], and that of the unweighted UniFrac distance was 0.95 [0.90; 0.98] (see Supplementary Figure 2), thus also indicative of their high predictive value. The difference between the 2 AUROCs was not significant (p=0.10).

**Figure 3.**
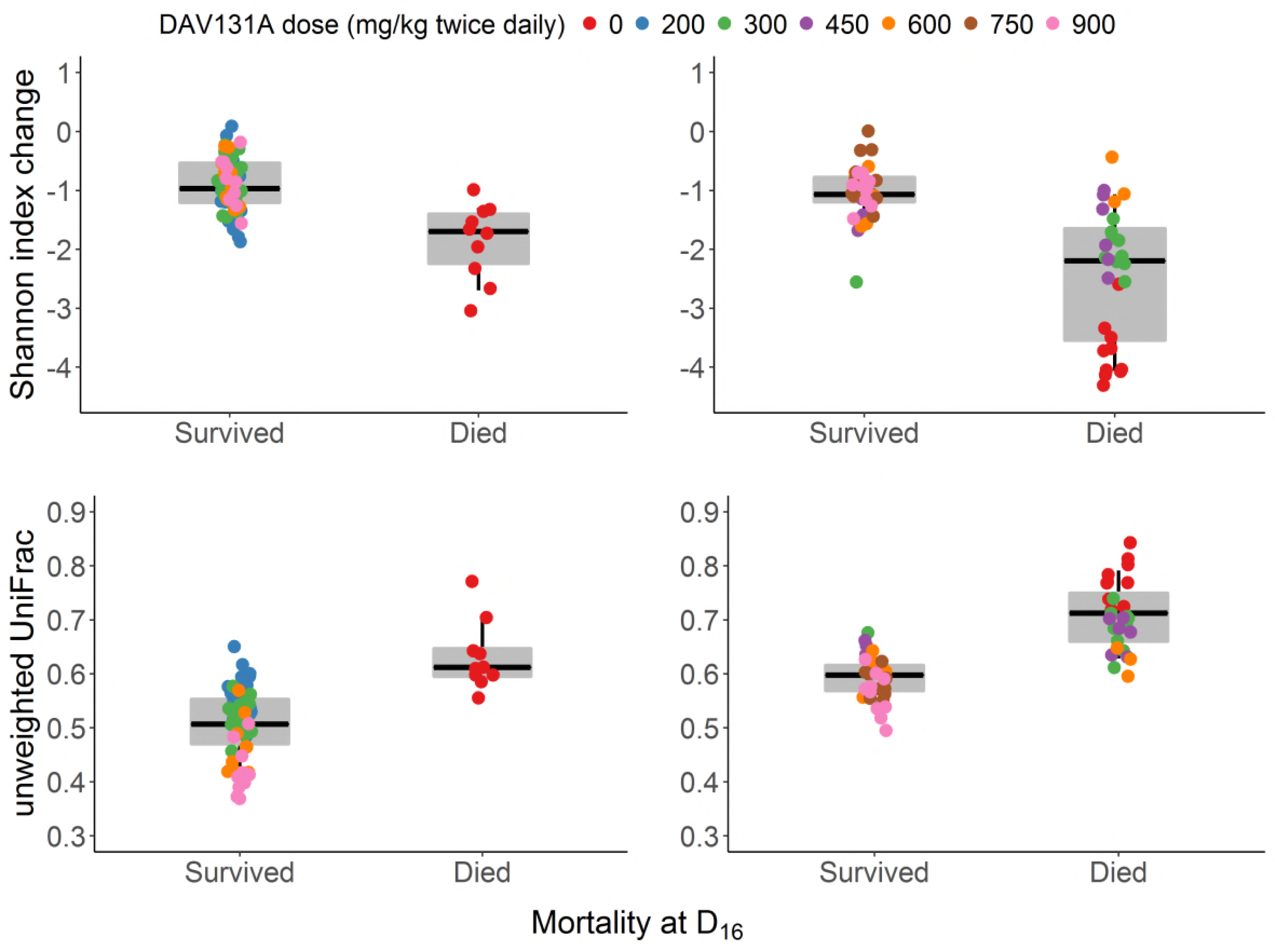
Change of Shannon index (top panel) and unweighted UniFrac distance (bottom panel) between D_0_ and D_3_ according to the occurrence of death by D_16_ in the moxifloxacin (left panel) or clindamycin (right panel) study. The boxes present the 25^th^ and 75^th^ percentiles and the horizontal black bar repo n value, while whiskers report 5^th^ and 95^th^ percentiles.

**Figure 4.**
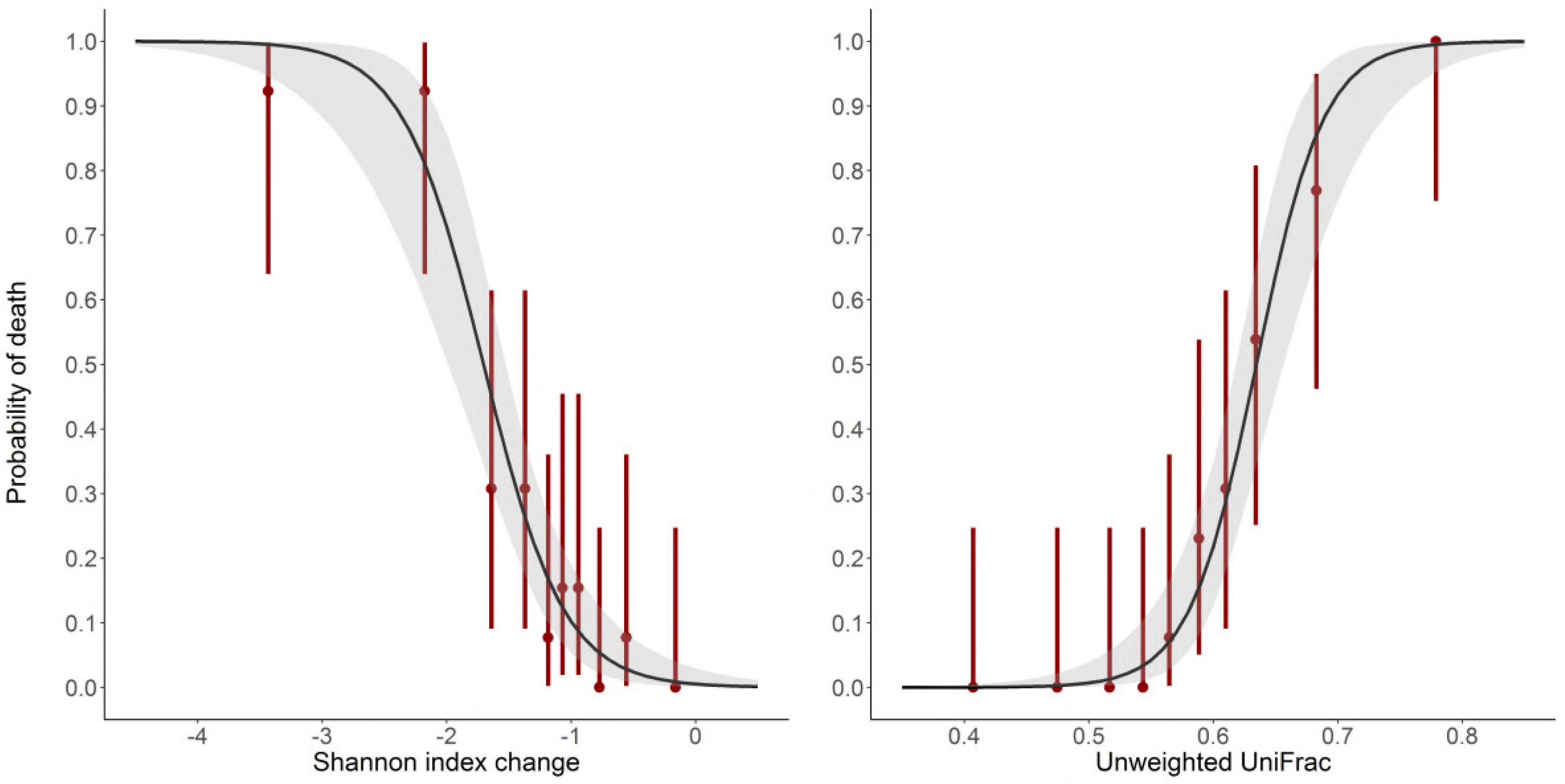
Logistic models of mortality according to the change of Shannon index (left panel, p<10^−15^) and unweighted UniFrac distance (right panel, p<10^−15^) between D_0_ and D_3_ after pooling data from antibiotic-treated animals in the moxifloxacin and clindamycin studies. Red bars represent the mortality rates and their 95% confidence intervals of deciles of the observed diversity indices. The shadded area present the 95% confidence interval of the predicted probability of death.

As these two indices were highly predictive of mortality, we further studied them by determining their optimal cut-off value best discriminating between death and survival by D_16_ using the Youden index. The value of the Shannon index change best discriminating between death and survival at D_16_ was −1.7 [−1.8; −1.2] (Supplementary Figure 3). The probability of observing a loss of diversity higher than −1.7 in hamsters which would die by D_16_ (sensitivity) was 0.71 [0.63; 0.95] and the probability of observing a loss of diversity lower than −1.7 in hamster which survived at D_16_ (specificity) was 0.96 [0.76; 0.99]. The best cut-off value of the unweighted UniFrac distance was 0.61 [0.58; 0.64] (Supplementary Figure 3). Associated sensitivity and specificity were 0.87 [95%CI, 0.79; 1.00] and 0.88 [0.72; 0.97], respectively. These values of sensitivity and specificity further illustrate the high predictability of these two diversity indices for the occurrence of the death of hamsters in these experiments.

Finally, in order to quantify the relationship between the loss of diversity and mortality, and to determine the maximal change of diversity required to limit the mortality rate to predefined values, we developed a logistic model of the probability of death according to the diversity observed in the intestinal microbiota. The model showed that small losses of diversity were sufficient to allow the development of severe colitis and death in a substantial number of animals. For instance, a reduction of the Shannon index between D_0_ and D_3_ by as little as 0.7 [95%CI, 0.4; 1.1] predicted to result in the death of 5% of the animals. The same mortality rate was predicted by an unweighted UniFrac distance of 0.51 [0.47; 0.55] between D_0_ and D_3._ Results for other mortality rates are presented in Table 3.

**Table 3.** Estimated change of Shannon index and unweighted UniFrac distances between D_0_ and D_3_ (and their 95% confidence intervals) needed to decrease the mortality rate to 50%, 10%, 5% and 1% in the hamster model of moxifloxacin-induced *Clostridium difficile* infection.

| Mortality rate | Change of Shannon index | unweighted UniFrac distance |
| --- | --- | --- |
| 10% | -1.0 [-1.2; -0.8] | 0.58 [0.55; 0.60] |
| 5% | -0.7 [-1.1; -0.4] | 0.55 [0.53; 0.58] |
| 1% | -0.2 [-0.7; 0.3] | 0.51 [0.47; 0.55] |

## Discussion

Our main result was the evidence of an association between the probability of hamster death and the antibiotic-induced loss of diversity of the intestinal microbiota at the time of *C. difficile* inoculation. Seemingly such a quantitative relationship had never been described. In this animal model, antibiotics perturb the structure and function of the intestinal microbiota, allowing the germination and growth of *C. difficile* spores, and the production of cytotoxic toxins leading to death of the animals (12). The protection provided by DAV131A through lowering the fecal concentration of active antibiotic, previously shown for moxifloxacin (22) was here extended to clindamycin, an antibiotic from a different class with a very different mode of action and spectrum of activity.

Our observations showed a clear relation between the loss of intestinal microbiota diversity and the development of *C. difficile* associated death in this model. Both α- and β-diversity indices studied had high predictive capacities for the ability of *C. difficile* spores to generate a lethal infection, the change of Shannon index between the beginning of antibiotic treatment and the time of *C. difficile* inoculation (for α-diversity) and the unweighted UniFrac distance (for β-diversity) appearing as the most predictive metrics. A link between the reduction of intestinal microbiota diversity after treatment with the glycylcycline antibiotic tigecycline has previously been reported in mice (24) but the precise quantitative relationship was not analyzed. In humans, the occurrence of *C. difficile* infection has been associated with a reduced diversity within the intestinal microbiota (25, 26). Here, we extend these observations to hamsters treated with either of two very different antibiotics moxifloxacin and clindamycin. We showed using various metrics, that individual-specific loss of diversity within the intestinal microbiota induced by antibiotics prior to *C. difficile* inoculation was highly predictive of the animals’ susceptibility to *C. difficile* infection, thus providing further insight into *C. difficile* pathophysiology. Furthermore, we were able to quantify this link, with even a small loss of diversity significantly increasing the risk of mortality. Indeed, a 0.7 reduction of the Shannon diversity index was associated with a 5% risk of death.

Furthermore, we observed that the loss of diversity was correlated to the concentration of active antibiotic in the fecal content. By adsorbing antibiotic residues reaching the colon after subcutaneous administration, DAV131A protected the microbiota against antibiotic-induced dysbiosis and reduced mortality in a dose dependent-manner. This approach appears to be promising as it might be extended to most classes of antibiotics, in addition to the two tested here, due to the wide adsorbing capacities of the product (21). Transposition to humans is currently ongoing. In a phase 1 clinical trial, DAV132, the human counterpart of DAV131A, was shown to reduce by more than 99% the fecal exposure to moxifloxacin in healthy volunteers, while the plasma concentration of the antibiotic remained unaffected; in subjects co-treated by moxifloxacin and DAV132, the diversity of the microbiota was protected from moxifloxacin-induced disruption (21). Further developments of this strategy to protect patients from the deleterious consequences of antibiotic treatments on the microbiota are currently ongoing.

## Material and Methods

### Hamster model of antibiotic-induced *C. difficile* infection

A previously developed hamster model of antibiotic-induced *C. difficile* infection was adapted to moxifloxacin (a fluoroquinolone antibiotic) and clindamycin (a lincosamide antibiotic) (27). After an 8-day acclimation period, male Golden Syrian hamsters (80-120 grams) received antibiotic by subcutaneous injection at a time designated as H_0_, once a day from day 1 (D_1_) to day 5 (D_5_). Administered doses were 30 mg/kg for moxifloxacin and 5 mg/kg for clindamycin. These doses were chosen as the lowest dose resulting in a 100% mortality rate in treated hamsters infected with *C. difficile* spores.

Animals were infected orally on day 3 (D_3_), 4 hours after antibiotic administration (H_4_), with 10^4^ spores of the non-epidemic *C. difficile* strain UNT103-1 (VA-11, REA J strain), TcdA+, TcdB+, cdtB−, vancomycin MIC = 2 μg/mL, moxifloxacin MIC = 16 μg/mL, clindamycin MIC > 256 μg/mL, ceftriaxone MIC = 128 μg/mL, obtained from Dr. Curtis Donskey, Ohio VA Medical Centre.

Vital status of the animals was evaluated daily until the end of the study at D16. Animals judged in a moribund stated were euthanized. All surviving hamsters were euthanized at D16.

### Ethics statement

Animals were housed in conformity with NIH guidelines (28). All procedures were conducted at the University of North Texas Health Science Center in Fort Worth (Texas, USA) in accordance with Protocol IACUC-2016-0015 approved by the local Institutional Animal Care and Use Committee.

### DAV131A

DAV131A is an activated charcoal-based adsorbent with high adsorption capacity (29). It was administered to hamsters by oral gavage after mixing with 0.25% w/v Natrosol^®^ 250 Hydroxyethylcellulose. Hamsters from placebo groups received Natrosol^®^ alone.

### Study design

Two studies with rather similar design were performed each with one antibiotic, moxifloxacin or clindamycin, in order to assess the protection provided by DAV131A against lethal antibiotic-induced *C. difficile* infection. DAV31A was administered *bis in die* (bid) to hamsters for 8 days, at H0 and H5 on D1, then at H-4 and H1 from D2 to D8.

In the moxifloxacin study, 70 animals were treated with moxifloxacin and 10 animals were left untreated. Groups of 10 or 20 antibiotic-treated animals were constituted according to the DAV131A unit dose administered: DAV131A placebo (MXF/0, n=10), 200 mg/kg (MXF/200, n=20), 300 mg/kg (MXF/300, n=20), 600 mg/kg (MXF/600, n=10) or 900 mg/kg (MXF/900, n=10). The control group was not treated by antibiotic and received DAV131A placebo.

In the clindamycin study, 60 animals were treated with clindamycin and 10 were left untreated. Groups of 10 antibiotic-treated animals were constituted according to the DAV131A unit dose administered: DAV131A placebo (CLI/0, n=10), 300 mg/kg (CLI/300, n=10), 450 mg/kg (CLI/450, n=10), 600 mg/kg (CLI/600, n=10), 750 mg/kg (CLI/750, n=10) or 900 mg/kg (CLI/900, n=10). The control group was not treated by antibiotic and received DAV131A placebo.

### Sample collection

For each animal, 2 fecal samples were collected, at D0 and D3. On D0, the fecal sample comprised pellets emitted in the 12 hours preceding the first antibiotic administration. On D3, samples were constituted by pellets emitted in the 12 hours following the third antibiotic administration; this surrounds the time at which animals were challenged by gavage with *C. difficile* spores (at 4h after antibiotic administration). Coprophagy of hamsters was not controlled, as this is a natural behavior in rodents. Fecal samples were stored at −80°C until further analysis.

### Measure of antibiotic concentrations

Fecal concentrations of active antibiotic were determined on fecal samples collected at D0 and D3 by a microbiological bioassay. On the day of the assay, feces were weighted, homogenized in sterile saline, and debris were eliminated by centrifugation. Fecal active moxifloxacin concentrations were measured using *B. subtilis* ATCC 6633 after incubation at 37°C for 24 hours (30). Fecal concentrations of active clindamycin were measured using *M. luteus* ATCC 9341 after incubation at 37°C for 24 hours (31). Data below the limit of quantification were imputed to 0.

### 16S rRNA gene bacterial community profiling

Microbial DNA was extracted using an extraction protocol optimized at GenoScreen, partially based on commercially available extraction kits (QIAamp DNA stool Kit, Qiagen, Germany) with the addition of chemical and mechanical lysis steps.

The V3-V4 region of the 16S rRNA gene was then amplified using an optimized and standardized amplicon-library preparation protocol (Metabiote^®^, GenoScreen, Lille, France). Positive (Artificial Bacteria Community comprising 17 different bacteria, ABCv2) and negative (sterile water) controls were also included. Briefly, PCR reactions were performed using 5 ng of genomic DNA and in-house fusion barcoded primers (final concentrations of 0.2 μM), with an annealing temperature of 50°C for 30 cycles. PCR products were purified using Agencourt AMPure XP magnetic beads (Beckman Coulter, Brea, CA, USA), quantified according to GenoScreen ‘s protocol, and mixed in an equimolar amount. Sequencing was performed using 250-bp paired-end sequencing chemistry on the Illumina MiSeq platform (Illumina, San Diego, CA, USA) at GenoScreen.

Raw paired-end reads were then demultiplexed per sample and subjected to the following process:

(1) search and removal of both forward and reverse primer using CutAdapt, with no mismatches allowed in the primers sequences; (2) quality-filtering using the PRINSEQ-lite PERL script (32), by truncating bases at the 3′ end with Phred quality score <30; (3) paired-end read assembly using FLASH (33), with a minimum overlap of 30 bases and >97% overlap identity.

Taxonomic and diversity analysis were performed using the Metabiote Online v2.0 pipeline (GenoScreen, Lille, France) which is partially based on the QIIME software v1.9.1 (34). Following the steps of pre-processing, chimera sequences were detected and eliminated (in-house method based on the use of Usearch 6.1). Then, clustering of similar sequences (97% identity threshold for an affiliation at the genus level on the V3-V4 regions of the 16S rRNA gene) was performed with Uclust v1.2.22q (35) through an open-reference OTU picking process and complete-linkage method, finally creating groups of sequences or “Operationnal Taxonomic Units” (OTUs). An OTU cleaning step corresponding to the elimination of singletons was performed. For each OTU, the most abundant sequence was considered as the reference sequence and taxonomically compared to the Greengenes database, release 13_8 (www.greengenes.gov) by the RDP classifier method v2.2 (36).

Various diversity indices were computed using QIIME (34). α-diversity metrics included the Shannon diversity index, the number of observed OTUs and the Chao1 index. In order to study the evolution of the bacterial diversity after the beginning of antibiotic treatment, we computed for each animal the difference between the values of these indices at D_3_ and D_0_. For β-diversity metrics, we computed the unweighted and weighted UniFrac distances, as well as Bray-Curtis dissimilarity for each animal between the samples collected at D_3_ and D_0_.

### Statistical analysis

For each study, we compared mortality rates at D_16_ and diversity indices across groups using nonparametric Fisher exact or Kruskall-Wallis tests, as appropriate. Fecal active antibiotic concentrations were compared according to DAV131A unit dose in antibiotic-treated hamsters using the Kruskall-Wallis test. In case of significant difference, post-hoc comparisons of each of the antibiotic-treated groups to the control group were performed using non parametric Fisher exact or Wilcoxon test with Benjamini-Hochberg ‘s correction for multiple testing. The correlations between active moxifloxacin or clindamycin fecal concentrations and diversity indices were studied using the Spearman rank correlation coefficient among antibiotic-treated hamsters.

We then compared for each study the fecal active antibiotic concentrations or diversity indices at D_3_ according to the vital status at D_16_ in antibiotic-treated hamsters, using the non-parametric Wilcoxon test. The predictability of death by D_16_ of the fecal active antibiotic concentration and of each studied diversity index was evaluated using the area under the Receiving Operator Curve (ROC) curve (AUROC) and its 95% confidence interval, computed using 2000 paired-bootstrap replicates. In the context of the present work, the AUROC can be interpreted as the probability that the diversity index will correctly rank 2 randomly chosen animals, 1 which would die by D16, and 1 which would survive (37).

In order to further analyze the link between microbial diversity and mortality by D16, we pooled the data of the 2 studies and performed a logistic regression of mortality by D16 according to diversity index in all antibiotic-treated hamsters. Diversity indices studied were those with the best predictive capacity among α- and β-diversity indices. Predictability was estimated using the AUROC and its 95% confidence interval. AUROCs of the 2 indices were compared using 2000 paired-bootstrap replicates. The best cut-off value for discriminating between hamsters which died and which survived at D_16_ was determined as the value allowing the maximization of both sensitivity and specificity, using the Youden index (38) and its 95% confidence interval. In the frame of the present study, sensitivity represents the probability of change of diversity between D_0_ and D_3_ being higher than a cut-off value in hamsters who will die by D_16_, and specificity is the probability of the change of diversity being lower than a cut-off value in hamsters who will survive until D_16_. The Youden index is computed as sensitivity + specificity – 1, and ranges between −1 and 1. A logistic model was then used to determine the diversity index values required to reduce mortality to various rates ranging from 1% to 10%.

Data are presented as number of observations n (%) or median (min; max). All tests were 2-sided with a type-I error of 0.05. All analyses were performed using R software v3.2.2.

## References

1. Dethlefsen L, Relman DA. 2011. Incomplete recovery and individualized responses of the human distal gut microbiota to repeated antibiotic perturbation. Proc Natl Acad Sci U S A 108 Suppl 1:4554–61.

2. Lichtman JS, Ferreyra JA, Ng KM, Smits SA, Sonnenburg JL, Elias JE. 2016. Host-microbiota interactions in the pathogenesis of antibiotic-associated diseases. Cell Rep 14:1049–1061.

3. Perez-Cobas AE, Gosalbes MJ, Friedrichs A, Knecht H, Artacho A, Eismann K, Otto W, Rojo D, Bargiela R, von Bergen M, Neulinger SC, Daumer C, Heinsen FA, Latorre A, Barbas C, Seifert J, dos Santos VM, Ott SJ, Ferrer M, Moya A. 2013. Gut microbiota disturbance during antibiotic therapy: a multi-omic approach. Gut 62:1591–601.

4. Theriot CM, Bowman AA, Young VB. 2016. Antibiotic-induced alterations of the gut microbiota alter secondary bile acid production and allow for *Clostridium difficile* spore germination and outgrowth in the large intestine. mSphere 1.

5. Jernberg C, Lofmark S, Edlund C, Jansson JK. 2010. Long-term impacts of antibiotic exposure on the human intestinal microbiota. Microbiology 156:3216–23.

6. Bergogne-Berezin E. 2000. Treatment and prevention of antibiotic associated diarrhea. Int J Antimicrob Agents 16:521–6.

7. Theriot CM, Young VB. 2015. Interactions between the gastrointestinal microbiome and *Clostridium difficile*. Annu Rev Microbiol 69:445–61.

8. Dubberke ER, Olsen MA. 2012. Burden of *Clostridium difficile* on the healthcare system. Clin Infect Dis 55 Suppl 2:S88–92.

9. Slimings C, Riley TV. 2014. Antibiotics and hospital-acquired *Clostridium difficile* infection: update of systematic review and meta-analysis. J Antimicrob Chemother 69:881–91.

10. Lessa FC, Mu Y, Bamberg WM, Beldavs ZG, Dumyati GK, Dunn JR, Farley MM, Holzbauer SM, Meek JI, Phipps EC, Wilson LE, Winston LG, Cohen JA, Limbago BM, Fridkin SK, Gerding DN, McDonald LC. 2015. Burden of *Clostridium difficile* infection in the United States. N Engl J Med 372:825–34.

11. CDC. 2017. Biggest Threats. https://www.cdc.gov/drugresistance/biggest_threats.html. Accessed May, 4th 2018

12. Best EL, Freeman J, Wilcox MH. 2012. Models for the study of *Clostridium difficile* infection. Gut Microbes 3:145–67.

13. Wilson KH, Silva J, Fekety FR. 1981. Suppression of *Clostridium difficile* by normal hamster cecal flora and prevention of antibiotic-associated cecitis. Infect Immun 34:626–8.

14. Larson HE, Borriello SP. 1990. Quantitative study of antibiotic-induced susceptibility to *Clostridium difficile* enterocecitis in hamsters. Antimicrob Agents Chemother 34:1348–53.

15. Elmer GW, Vega R, Mohutsky MA, McFarland LV. 1999. Variable time of onset of *Clostridium difficile* disease initiated by antimicrobial treatment in hamsters. Microbial Ecology in Health and Disease 11:163–168.

16. Metzker ML. 2010. Sequencing technologies - the next generation. Nat Rev Genet 11:31–46.

17. Shannon C. 1948. A mathematical theory of communication. The Bell System Technical Journal 27:623–56.

18. Chao A. 1984. Nonparametric estimation of the number of classes in a population. Scandinavian Journal of Statistics 11:265–70.

19. Lozupone C, Knight R. 2005. UniFrac: a new phylogenetic method for comparing microbial communities. Applied and Environmental Microbiology 71:8228–35.

20. Bray J, Curtis J. 1957. An ordination of the upland forest communities of southern Wisconsin. Ecological Monographs 27:325–49.

21. de Gunzburg J, Ghozlane A, Ducher A, Le Chatelier E, Duval X, Ruppé E, Armand-Lefèvre L, Sablier-Gallis F, Burdet C, Alavoine L, Chachaty E, Augustin V, Varastet M, Levenez F, Kennedy S, Pons N, Mentré F, Andremont A. 2018. Protection of the human gut microbiome from antibiotics. J Infect Dis 217:628–636.

22. Burdet C, Sayah-Jeanne S, Nguyen TT, Miossec C, Saint-Lu N, Pulse M, Weiss W, Andremont A, Mentré F, de Gunzburg J. 2017. Protection of hamsters from mortality by reducing fecal moxifloxacin concentration with DAV131A in a model of moxifloxacin-induced *Clostridium difficile* colitis. Antimicrob Agents Chemother 61.

23. Hosmer D, Lemeshow S. 2000. Applied Logistic Regression, USA.

24. Bassis CM, Theriot CM, Young VB. 2014. Alteration of the murine gastrointestinal microbiota by tigecycline leads to increased susceptibility to *Clostridium difficile* infection. Antimicrob Agents Chemother 58:2767–74.

25. Schubert AM, Rogers MA, Ring C, Mogle J, Petrosino JP, Young VB, Aronoff DM, Schloss PD. 2014. Microbiome data distinguish patients with *Clostridium difficile* infection and non-*C. difficile*-associated diarrhea from healthy controls. MBio 5:e01021–14.

26. Zhang L, Dong D, Jiang C, Li Z, Wang X, Peng Y. 2015. Insight into alteration of gut microbiota in *Clostridium difficile* infection and asymptomatic *C. difficile* colonization. Anaerobe 34:1–7.

27. Phillips ST, Nagaro K, Sambol SP, Johnson S, Gerding DN. 2011. Susceptibility of hamsters to infection by historic and epidemic BI *Clostridium difficile* strains during daily administration of three fluoroquinolones. Anaerobe 17:166–9.

28. National Research Council. 2011. Guide for the care and use of laboratory animals. The National Academy Press, Washington D.C., USA.

29. Grall N, Massias L, Nguyen TT, Sayah-Jeanne S, Ducrot N, Chachaty E, de Gunzburg J, Andremont A. 2013. Oral DAV131, a charcoal-based adsorbent, inhibits intestinal colonization by beta-lactam-resistant *Klebsiella pneumoniae* in cefotaxime-treated mice. Antimicrob Agents Chemother 57:5423–5.

30. Kampougeris G, Antoniadou A, Kavouklis E, Chryssouli Z, Giamarellou H. 2005. Penetration of moxifloxacin into the human aqueous humour after oral administration. The British Journal of Ophthalmology 89:628–31.

31. Courvalin O, Leclercq R, Rice L. 2010. Antibiogram. ASM Press.

32. Schmieder R, Edwards R. 2011. Quality control and preprocessing of metagenomic datasets. Bioinformatics 27:863–4.

33. Magoc T, Salzberg SL. 2011. FLASH: fast length adjustment of short reads to improve genome assemblies. Bioinformatics 27:2957–63.

34. Caporaso JG, Kuczynski J, Stombaugh J, Bittinger K, Bushman FD, Costello EK, Fierer N, Pena AG, Goodrich JK, Gordon JI, Huttley GA, Kelley ST, Knights D, Koenig JE, Ley RE, Lozupone CA, McDonald D, Muegge BD, Pirrung M, Reeder J, Sevinsky JR, Turnbaugh PJ, Walters WA, Widmann J, Yatsunenko T, Zaneveld J, Knight R. 2010. QIIME allows analysis of high-throughput community sequencing data. Nat Methods 7:335–6.

35. Edgar RC. 2010. Search and clustering orders of magnitude faster than BLAST. Bioinformatics 26:2460–1.

36. Wang Q, Garrity GM, Tiedje JM, Cole JR. 2007. Naive Bayesian classifier for rapid assignment of rRNA sequences into the new bacterial taxonomy. Appl Environ Microbiol 73:5261–7.

37. Fawcett T. 2006. An introduction to ROC analysis. Pattern Recognition Letters 27:861–74.

38. Youden WJ. 1950. Index for rating diagnostic tests. Cancer 3:32–5.

